# Rank-dependent social inheritance determines social network structure in a wild mammal population

**DOI:** 10.1101/2020.04.10.036087

**Authors:** Amiyaal Ilany, Kay E. Holekamp, Erol Akçay

## Abstract

The structure of animal social networks influences survival and reproductive success, as well as pathogen and information transmission. However, the general mechanisms determining social structure remain unclear. Using data on 73,767 social interactions among wild spotted hyenas over 27 years, we show that a process of social inheritance determines how offspring relationships are formed and maintained. The relationships of offspring with other hyenas are similar to those of their mothers over up to six years, and the degree of similarity increases with maternal social rank. The strength of mother-offspring relationship affects social inheritance and is positively correlated with offspring longevity. These results confirm the hypothesis that social inheritance of relationships can structure animal social networks and be subject to adaptive tradeoffs.

## 1 Introduction

The fine scale social structure of animal populations plays an important role in all social processes, including pathogen and cultural transmission^1,2,3,4^ and the evolution of social behaviors^5,6^. For these reasons, social structure and an individual’s position in it, affects reproductive success, longevity, and sexual selection^7,8,9^. An important aspect of social structure is the varying associations between different individuals, as represented by social networks. Research in the last few decades has started to elucidate patterns in social networks across animal species. These studies have been mostly descriptive (with some prominent exceptions such as the model of Seyfarth^10^), but a new generation of studies are trying to explain observed patterns using generative models^11,12,13,14^. In one of these studies, Ilany and Akçay recently proposed social inheritance, defined as a tendency of offspring social affiliations to resemble those of their parents, as a general process that can explain the structure of social networks of multiple species^12^. They showed that the structure of model networks where offspring tend to inherit (via passive or active copying) their parents’ social affiliations is similar to those of observed populations^12^. Most importantly, social inheritance of maternal associations leads to clustering, a key feature of social networks and distinguishes them from other types of networks^15^. As such, social inheritance may be crucial to the maintenance of stability in social networks.

Social inheritance is already empirically demonstrated for some aspects of social position: individuals in many species socially inherit maternal dominance ranks, which determine priority of access to resources and are calculated from observed agonistic interactions^16,17,18^. The inheritance of rank is likely to be non-genetic, because rank shows high plasticity in response to social and environmental factors^17,18^. In rhesus macaques^19^ as well as in African elephants^20^ the offspring of socially affiliated mothers are also likely to be affiliated. More generally, evidence from primates suggests that mothers may influence the development of their offspring’s social ties both passively and actively^21,22,23^. In African elephants, the network position (betweenness) of mothers was the strongest predictor of their daughters’ position a decade later, despite the population experiencing stress from poaching and a drought^20^.

These findings provide strong indirect evidence that inheritance of social relationships plays an important role in many species. In this study we directly demonstrate social inheritance in the spotted hyena (*Crocuta crocuta*), using data from 27 years of continuous field observations. Spotted hyenas live in stable groups, called clans, that are far more complex than those of other mammalian carnivores, resembling the societies of Old World primates such as baboons or macaques in their size and structure^24^. The size of hyena clans depends on local prey abundance, and may vary from only a few individuals to more than a hundred^24^. Hyena clans usually contain several matrilineal kin groups spanning multiple generations, with low average relatedness among clan members^25^. Wild spotted hyenas live up to 26 years. They can discriminate both maternal and paternal kin from unrelated hyenas^26,27^. They contend with their clan mates for access to killed prey, but high-ranking individuals maintain priority of access to food^24^. The long-term social network dynamics of hyenas are determined by a complex set of factors, including environmental effects such as rainfall and prey availability, individual traits such as sex and social rank, and also structural effects such as the tendency to close triads and form bonds with highly connected individuals^28^.

Using our long-term dataset of spotted hyena social interactions, we find that offspring social associations with individuals in their social group resemble their mothers’ associations with those same individuals, providing direct evidence for social inheritance. Importantly, social inheritance remains strong up to six years after leaving the communal den, even though the mother-offspring association itself decreases in strength. We also show that the strength of social inheritance is positively correlated with mother-offspring social bond and maternal rank. Finally, we show that social inheritance is positively correlated with both offspring and mother survival, even after controlling for maternal rank, a known factor affecting offspring survival. These results provide evidence that social inheritance is an important adaptation shaping animal social networks.

## 2 Results and Discussion

We quantified social networks using yearly association indices (AIs), defined as the number of times two individuals were observed together in a given year divided by the total number of times either were observed. We then quantify the similarity between two individuals’ social connections in a given year by looking at the correlation of their association indices with all other individuals (Fig. 1A). Social inheritance should result in a positive correlation between the association indices of mother and offspring with all other individuals in a population. Therefore, we first measured the mother-offspring correlation in AIs with others, and compared that to correlations between all other pairs of individuals (Fig. 1B; Fig. 1C). We found that the social associations of offspring were similar to those of their mothers, in contrast to a much weaker correlation within other pairs of hyenas (LMM, *β* = −0.32 ± 0.02, *P* < 0.001). Moreover, associations of mothers before their offspring left the den also predict those of their offspring in the following year (Fig. SI-5), which suggests that offspring indeed inherit their mothers’ existing connections, rather than mothers acquiring offspring connections or mothers and offspring forming new social ties together.

**Figure 1:**
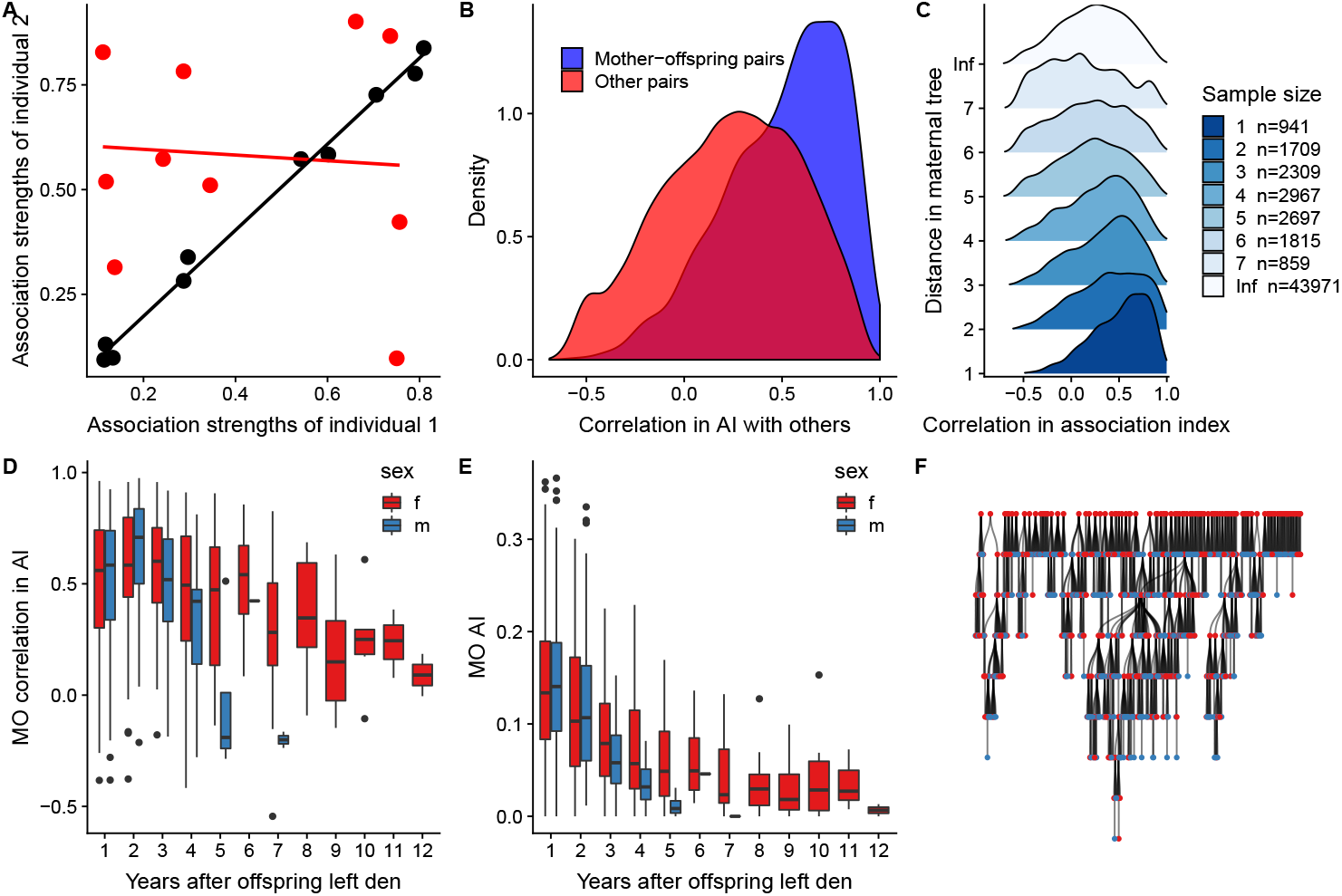
Social inheritance and its ontogeny in spotted hyenas. **A)** An illustration of correlation in association strengths, our measure of social inheritance. Association index measures the strength of association between two hyenas over one calendar year. Correlation in association index measures the similarity in association with others between two hyenas over one calendar year. In this toy example black points demonstrate a pair of individuals who are similar in their associations with other, whereas red points represent a pair whose associations with others are dissimilar. **B)** A comparison of densities of correlations in AIs within pairs of hyenas: mother-offspring pairs vs. other pairs demonstrating that mother-offpsring pairs have higher correlations than other pairs. **C)** A comparison of densities of correlations in AIs within pairs of hyenas, as a function of the within-pair distance in the maternal pedigree. Sample size is the number of dyads for each distance on the tree. **D)** Ontogeny of social inheritance. Boxplots depict the distribution of correlation between the AIs of mothers and those of their offspring, starting with the first year in which the offspring was observed at least 20 times away from the den. **E)** Ontogeny of mother-offspring relationship. Boxplots depict the distribution of AIs of mothers and their offspring, starting with the first year the offspring was observed at least 20 times away from the den. **F)** The hyena maternal pedigree. This tree shows all known maternal relationships within the Talek clan of spotted hyenas over 27 years. Blue and red circles depict females and males, respectively. N = 1320 hyenas.

Looking at the ontogeny of the mother-offspring correlation in AIs, we found that in the first four (for males) or six (for females) years after the offspring left the den, its social relationships remained similar to its mother’s (Fig. 1D, Table 1). The median correlation coefficient of mother vs. offspring AIs with other hyenas varied between 0.44 and 0.67 in the first six years in which they overlapped. This similarity in social relationships remained high even when the strength of relationship between the offspring and its mother decreased over the years (Fig. 1E, Table SI-1), from a median of 0.14 in the first year after the offspring left the den, to 0.05 in the sixth year. These results show that although social inheritance may initially depend on close association between the mothers and their offspring, it remains stable even after the mother-offspring association has subsided.

**Table 1:** Ontogeny of social inheritance. Mixed model outcome for the correlation between mother and offspring association with other hyenas as a function of offspring sex and years since leaving den. Data include the first six years after offspring left the den. Mother-offspring pair ID was set as a random factor. *N* = 532.

|  | Estimate | SE | df | t value | P |
| --- | --- | --- | --- | --- | --- |
| (Intercept) | 0.56 | 0.03 | 504.07 | 16.13 | 0.00 |
| years since leaving den | -0.02 | 0.01 | 467.59 | -1.82 | 0.07 |
| offspring sex (ref:f) | 0.06 | 0.05 | 514.72 | 1.28 | 0.20 |
| (years since leaving den):(offspring sex) | -0.04 | 0.02 | 462.77 | -2.04 | 0.04 |

In spotted hyenas, social rank plays a major role in structuring the clan, with important consequences for fitness^24^. Rank may affect social inheritance in at least two non-mutually exclusive ways: first, offspring of higher-ranked individuals are expected to face fewer constraints in choosing social partners than lower-ranked offspring, both due to having more time for socializing, and also presumably to having more willing partners^28^. Second, offspring of lower-ranked individuals may benefit from forming different associations than their parents to compensate for their low rank. Both the constraint and compensation hypotheses predict a weaker mother-offspring correlation in association indices for lower-ranked mothers. As Fig. 2A-E show, this prediction is confirmed by our data, but only for offspring after their first year out of the den (Tables SI-5,SI-6). In the first year most mother-offspring pairs have high correlation of association indices, regardless of rank. Interestingly, while the mean mother-offspring correlation declines with lower rank, the variability of correlations increases, which means some parents and offspring are able to maintain similar connections while others cannot. This suggests that social constraints faced by lower-ranked individuals may play a role in the decline of mother-offspring correlation over time. On the other hand, offspring of low-ranked mothers tended to form stronger associations overall with other hyenas than their mothers do (Fig. 2F; linear mixed model with mother ID and year set as random factors: *β* = 0.0007 ± 0.0001, *P* < 0.001), which suggests that offspring may compensate for their low rank by socializing more. We also find that after controlling for maternal rank, mother-offspring association strength, and offspring sex, offspring were more likely to inherit maternal associations if there were more close relatives in the clan (SI-3), but this effect disappeared when considering also more distant relatives (SI-2).

**Figure 2:**
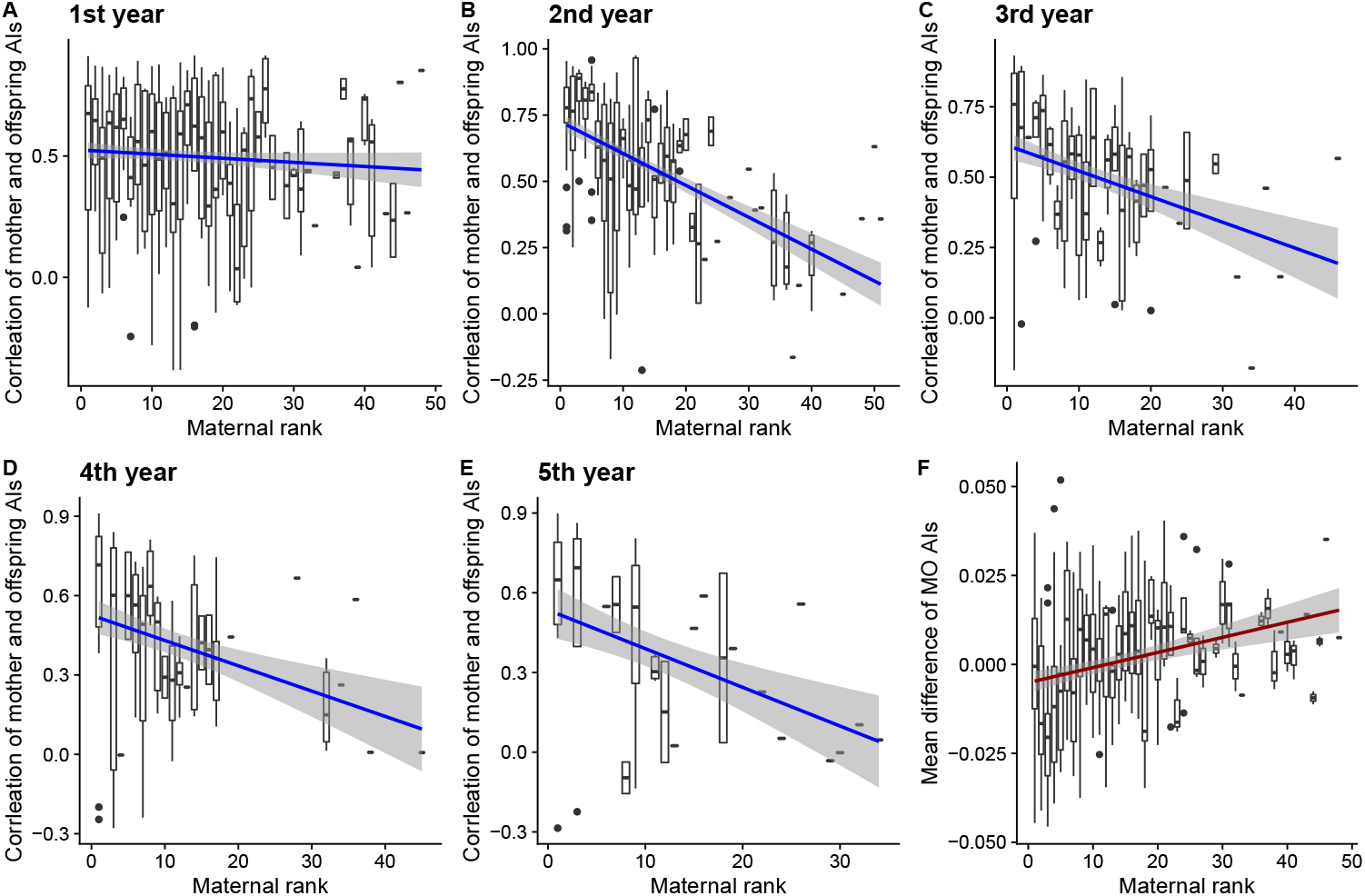
The effect of maternal social rank on social inheritance. **A-E)** Boxplots depict mother-offspring correlation in association indices (AIs) for each maternal rank. Year notes time since offspring left the den. Total *N* = 532. **F)** The effect of maternal rank on deviation of offspring associations from those of their mothers. Boxplots depict the mean differences between the AIs of offspring and those of their mothers in the first year after leaving the den (*N* = 342). By convention, smaller numbers represent higher ranks.

Next, we document that social inheritance is associated with longevity of both mothers and female offspring. On average, offspring who formed social associations more similar to those of their mothers in their first year of overlap survived longer (Fig. 3A). For offspring of alpha females (highest-ranked), no social inheritance (a mother-offspring correlation of zero) would translate to a predicted increase of 528% in hazard compared to offspring with maximum social inheritance (mother-offspring correlation of 1). In contrast, for an offspring of a mother ranked 30th in the clan, no social inheritance will translate to a predicted decrease of 65% in hazard. The effect of social inheritance on offspring longevity held even when controlling for maternal rank, a known predictor of longevity^29^. However, since social inheritance is also correlated with mother-offspring association strength in the first year (Fig. SI-4; linear mixed model with mother and year as random effects: *β* = 0.31 ± 0.02), it is possible that the association link between social inheritance and longevity is not causal, but instead that both are caused by increased mother-offspring association. Indeed, structural equation modeling revealed that the strength of mother-offspring association in the first year after leaving the den predicts both off-spring longevity and social inheritance (standardized coefficients in the best supported model: mother-offspring correlation in association on association strength: *β* = 0.67; offspring’s last age on association strength: *β* = 0.41; Table SI-4).

**Figure 3:**
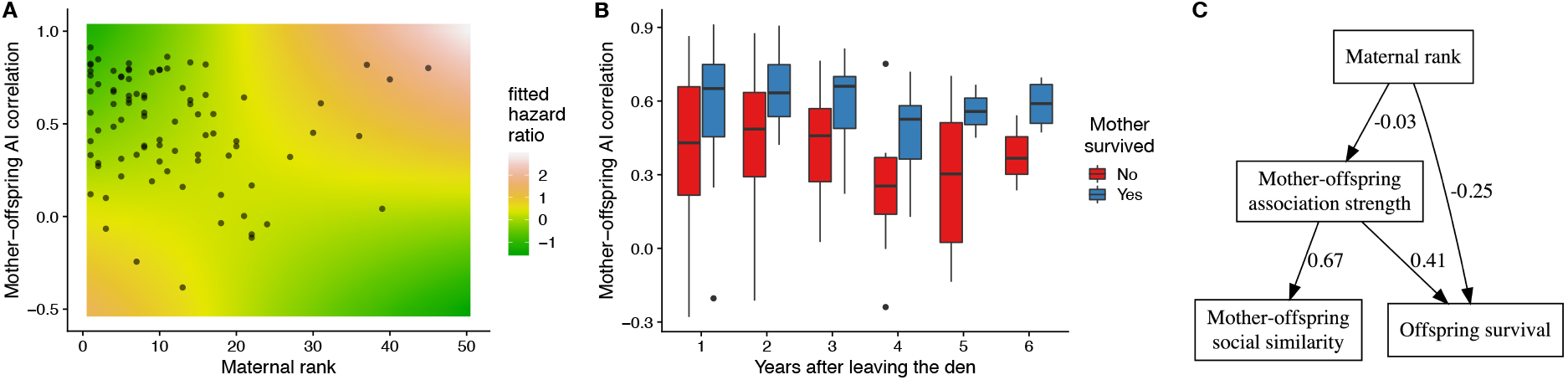
Social inheritance is associated with both offspring and mother survival. **A)** Fitted hazard ratios from a model of offspring survival depict dependence on maternal rank and mother-offspring AI correlation (higher hazard ratios reflect lower survival chances). Points indicate observed values (*N* = 88). **B)** Social inheritance predicts maternal survival. The strength of social inheritance (correlation of mother and offspring association indexes with other hyenas) was lower for mothers that did not survive to the following year (red boxplots), compared to those who did survive (blue). This trend was consistent across offspring ages. *N* = 206. **C)** Mother-offspring association predicts both social inheritance and offspring survival. Path diagram showing correlations between maternal rank, mother-offspring association, social inheritance, and offspring survival. The partial regression coefficients for each arrow are given.

Finally, our data suggest that social inheritance by offspring is associated with higher survivorship of mothers. Mothers of offspring who were more similar to them in social associations were more likely to survive to the following year (Fig. 3B; likelihood ratio test = 17.35, *d.f.* = 1, *P* < 0.0001; logistic regression of maternal survival with maternal rank and offspring age as fixed effects). Offspring that do not socialize with the associates of their mothers may thus provide a cue that those mothers are aging.

Taken together, our results suggest that social inheritance plays an important role in structuring hyena social networks. This provides further support for Ilany and Akçay’s hypothesis that in species with stable social groups, the inheritance of social connections from parents is the cornerstone of social structure. Furthermore, we show that in a social carnivore, social relationships, and the position within the social networks they represent, are socially inherited similarly to how social rank is inherited in this species^17^. Similar patterns were found in African elephants^20^ and in some primate species^23^, implying that social inheritance of associations could be common and adaptive. We quantified social inheritance by measuring the similarity of social connections of mothers and their offspring to third individuals. This measure of social similarity adds a new dimension to the analysis of animal societies. Whereas the strength of relationship among two individual is widely used, social similarity of two individuals can provide additional information about a relationship, its origins, and consequences.

In spotted hyenas, females usually dominate males as a result of higher social support^30^. Our results suggest a bi-directional feedback between social relationships and social rank; dominance affects social inheritance, whereas the resulting social relationships affect social hierarchy^31^. Social rank and network position affect one another, and also influence survival probabilities. The differential effect of both sex and maternal rank on social inheritance suggests that hyenas actively modify their social associations to accommodate multiple environmental and social factors. Offspring of low-ranked mothers suffer the consequences of inheriting a low rank, including limited access to prey compared to their high-ranked conspecifics. The social associations of these offspring diverge the most from those of their mothers, suggesting that they are either constrained from inheriting their mother’s associations or attempt to compensate for their mother’s low rank. Our data provides support for both options.

A challenge lies in finding the extent to which copying of maternal associations is active or passive and how much is it driven by the offspring (inheritance) or the parent (bequeathal). There is evidence that mothers do actively influence their offspring’s social connections^21,22,23^. However, the extent and the precise mechanisms by which mothers can bias their offspring’s social networks remain to be worked out. Another open question is whether there is genetic variability in the propensity of inheriting or bequeathing social connections, and how selection acts on such genetic variation. The association between social inheritance and survival we document suggests that active social inheritance may be selected for, even if social inheritance is not causal to survival. Overall, our results highlight the role social inheritance plays at the nexus of social network structure and life history.

## Materials and Methods

### Data collection

We used daily observations of the Talek clan, conducted between 1989 and 2015, in the Masai Mara National Reserve, Kenya^32^. This clan occupies a group territory of roughly 70 km^2^. Hyenas are individually recognized by their unique spots. Sex is determined by the morphology of the erect phallus. The position of an individual in a matrix ordered by submissive behavior displayed during agonistic encounters determines its social rank^24^. To estimate the effect of social inheritance on survival, we recorded the last known age of each female offspring. We used a conservative approach, including only females who were dead by the end of the study (*N* = 88). Male survival is difficult to estimate, as males disperse and their age at death is often unknown.

### Constructing social networks

We could individually identify all hyenas belonging to the clan. We removed from the analysis transient individuals, such as those visiting the Talek territory to hunt for a day. Hyenas were included in the analysis only if they were observed at least 20 times in at least one year, and were considered absent in a given year if observed < 20 times during that year. Hyenas that were included were observed an average ± SE of 98.8 ± 1.9 times per year. Cubs, defined as young hyenas that were still residing at dens^24^, were not included. Observations at dens were also excluded to eliminate bias towards lactating females. Thus, the 1st year of offspring after leaving the den was defined as the 1st year in which it was observed at least 20 times in non-den sessions. Hyenas were assigned to a single observation session if they were found together, separated from other hyenas by at least 200 m, but usually at least 1 km^33^. From the observations we calculated, for each pair per year, the simple-ratio association index, that describes the strength of an association on a range from zero to one^34^.

### Social similarity

We define social similarity between two individuals as the Pearson correlation coefficient of their association indexes with all other individuals in the network. Thus, two individuals will have high social similarity if their connections with other individuals are of similar magnitude.

### Maternal pedigree

All natal clan members were observed during their first three months of life. The identity of mothers was determined by suckling behavior. From these identities we constructed a maternal pedigree, where we could measure for each pair of hyenas the distance as the path length (number of steps) between individual nodes on the pedigree (Fig. 1F). For example, the distance between two brothers is two: one step of brother A to its mother, and a second step from the mother to brother B. Hyenas that were not assigned a mother (such as immigrant males) had infinite distance to other hyenas on this tree.

### Survival Analysis

We used Cox proportional hazards models to determine how offspring risk of death was affected by the interaction of social inheritance and maternal rank. For each mother-offspring pair we used only social inheritance from the first year after the offspring had left the den. We used the ‘survival’ R package (version 3.1-7).

### Statistical Analysis

The factors affecting ontogeny of social inheritance and ontogeny of mother-offspring relationships were tested using generalized linear mixed models (GLMM). Similarly, GLMMs were used to test the impact of maternal rank on social inheritance. Walds z test and likelihood ratio tests (LRTs) were applied to determine the significance of fixed and random effects^35^. These analyses used the lmerTest R package (version 3.1-0). Logistic regression was used to test the impact of social inheritance on maternal survival to the following year.

### Structural Equation Modeling

Structural equation models unite multiple predictor and response variables in a single causal network^36^. We tested the causal relationships between maternal rank, mother-offspring association strength (MOAS), the correlation of mother and offspring association with others (MOC), and offspring last known age. Only female offspring were used, because males disperse and thus their longevity is difficult to estimate. We tested a series of plausible relationships between MOC, MOAS, maternal rank and offspring longevity, and used model selection to assess fit to data. To account for the data structure we used piecewise SEMs^36^, in which mother identity was set as a random effect in all models. Table SI-4 presents the tested relationships.

## Supplementary Tables and Figures

**Table SI-1:**
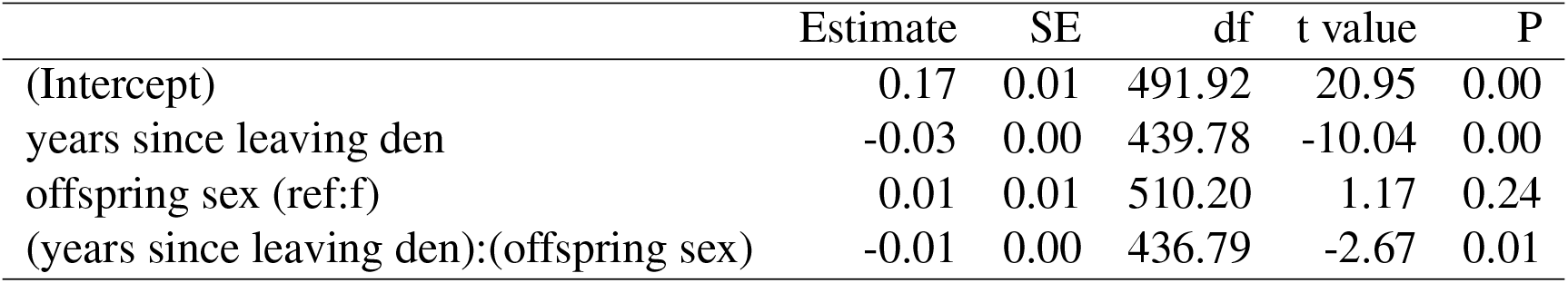
Ontogeny of mother-offspring relationship. Mixed model outcome for the association strength between mother and offspring as a function of offspring sex and years since leaving den. Data include the first six years after offspring left the den. Mother-offspring pair ID was set as a random factor. *N* = 532.

|  | Estimate | SE | df | t value | P |
| --- | --- | --- | --- | --- | --- |
| (Intercept) | 0.17 | 0.01 | 491.92 | 20.95 | 0.00 |
| years since leaving den | -0.03 | 0.00 | 439.78 | -10.04 | 0.00 |
| offspring sex (ref:f) | 0.01 | 0.01 | 510.20 | 1.17 | 0.24 |
| (years since leaving den):(offspring sex) | -0.01 | 0.00 | 436.79 | -2.67 | 0.01 |

**Table SI-2:**
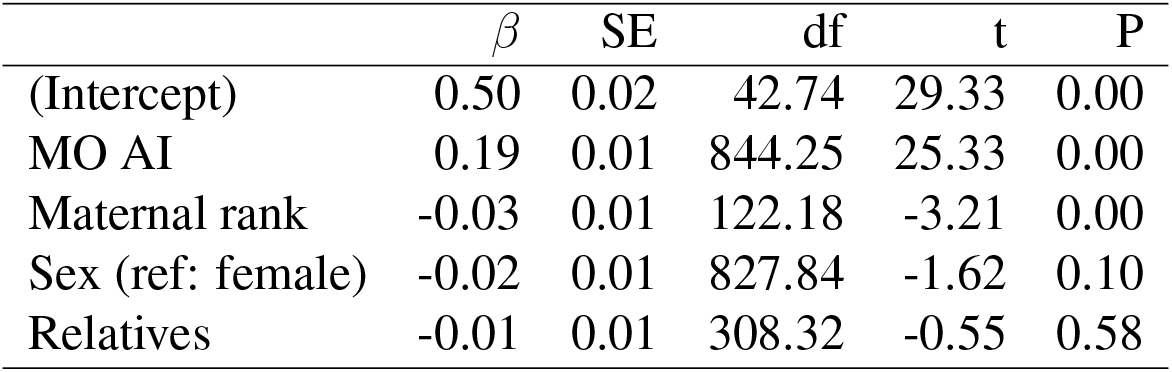
Factors affecting social inheritance considering close and distant relatives. Model outcome for the correlation between mother and offspring association with other hyenas. MO AI stands for the mother-offspring association index. Relatives are the number of present hyenas that are less than 5 steps away in the maternal pedigree.

**Table SI-3:** Factors affecting social inheritance considering close relatives. Model outcome for the correlation between mother and offspring association with other hyenas. MO AI stands for the mother-offspring association index. Relatives are the number of present hyenas that are less than 3 steps away in the maternal pedigree.

| | $\beta$ | SE | df | t | P |
| --- | --- | --- | --- | --- | --- |
| (Intercept) | 0.51 | 0.02 | 48.74 | 27.12 | 0.00 |
| MO.AI | 0.19 | 0.01 | 1697.58 | 35.42 | 0.00 |
| Maternal rank) | -0.02 | 0.01 | 118.21 | -2.11 | 0.04 |
| Sex (ref: female) | -0.02 | 0.01 | 1683.34 | -2.35 | 0.02 |
| Relatives | 0.02 | 0.01 | 1014.57 | 2.71 | 0.01 |

**Table SI-4:** Model selection for structural equation modeling. The tested series of models include relationship between mother-offspring association strength (MOAS), correlation of mother and offspring association with others (MOC), maternal social rank (MRank), and off-spring last known age (Age). All models were linear mixed models, with mother identity set as a random effect.

| Models | Random effects | $AIC_c$ | Weight |
| --- | --- | --- | --- |
| MOC ~ MOAS | Mother | 30.10 | 0.57 |
| MOAS ~ MRank | Mother |  |  |
| Age ~ MOAS + MRank | Mother |  |  |
| MOC ~ MOAS | Mother | 30.67 | 0.43 |
| MOAS ~ MRank | Mother |  |  |
| Age ~ MOAS + MRank + MOC | Mother |  |  |

**Table SI-5:** Social inheritance and maternal rank in the 1st year. Mixed model outcome for the strength of social inheritance as a function of maternal rank, mother-offspring association strength, and offspring sex, in the 1st year after leaving den. Mother ID and year were set as random factors. *N* = 684.

|  | Estimate | Std. Error | df | t value | Pr(>\|t\|) |
| --- | --- | --- | --- | --- | --- |
| (Intercept) | 0.138 | 0.035 | 88.969 | 4.001 | 0.000 |
| mother-offspring AI | 2.531 | 0.106 | 642.091 | 23.951 | 0.000 |
| maternal rank | 0.001 | 0.001 | 95.778 | 0.432 | 0.667 |
| offspring sex (ref:f) | -0.013 | 0.013 | 656.698 | -0.974 | 0.330 |

**Table SI-6:** Social inheritance and maternal rank in subsequent years. Mixed model outcome for the strength of social inheritance as a function of maternal rank, mother-offspring association strength, and offspring sex, in subsequent years after the 1st year after leaving den. Mother-offspring pair ID and year were set as random factors. *N* = 1058.

|  | Estimate | Std. Error | df | t value | Pr(>\|t\|) |
| --- | --- | --- | --- | --- | --- |
| (Intercept) | 0.328 | 0.029 | 112.778 | 11.210 | 0.000 |
| mother-offspring AI | 2.898 | 0.094 | 989.605 | 30.857 | 0.000 |
| maternal rank | -0.005 | 0.001 | 127.285 | -4.445 | 0.000 |
| offspring sex (ref:f) | -0.013 | 0.021 | 179.169 | -0.638 | 0.524 |

**Figure SI-1:**
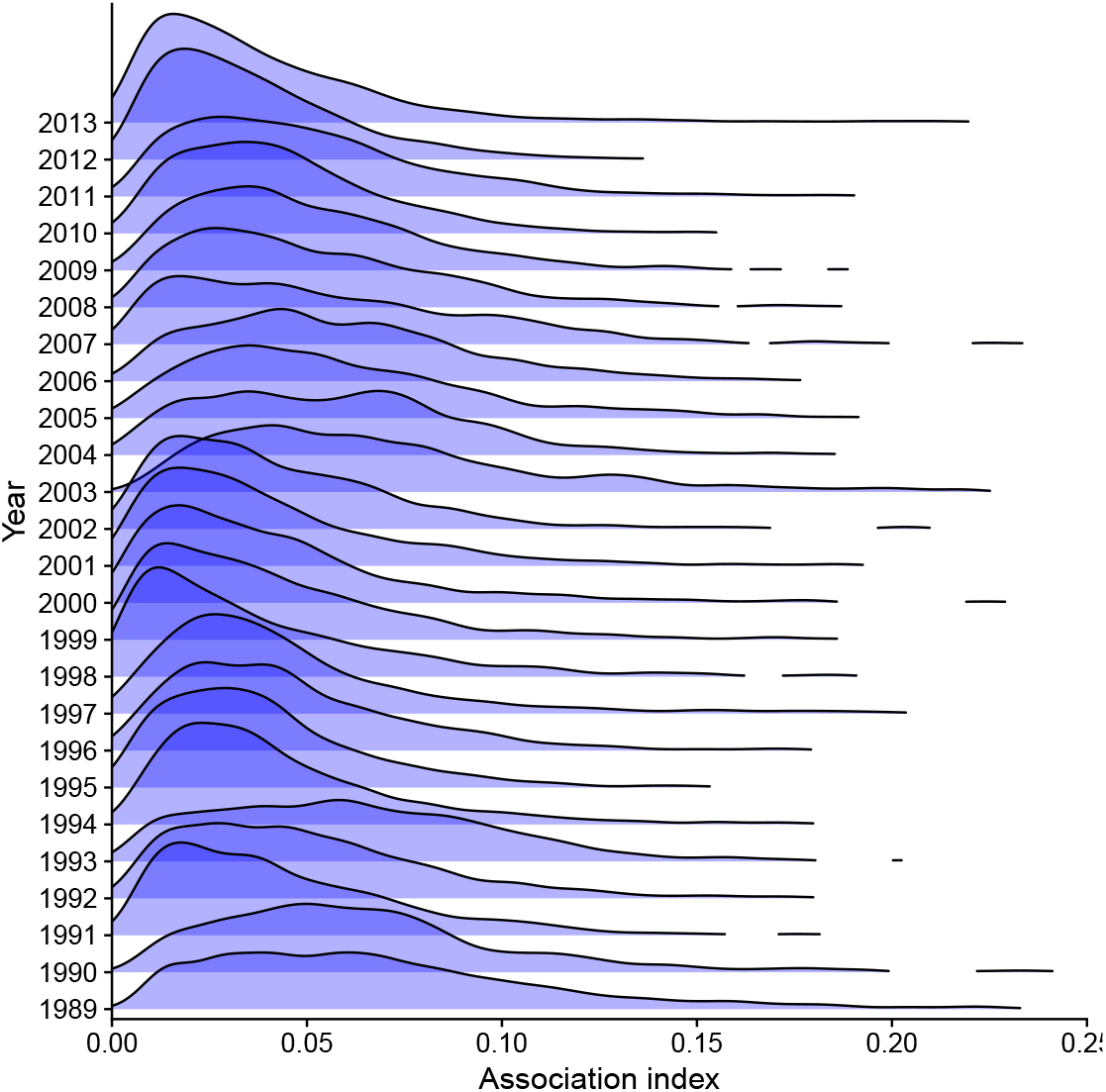
Distribution of association indices in spotted hyenas. Association index measures the strength of association between two hyenas over one calendar year. Shading is used to show overlapping distributions. Between 1989 and 2015, the distribution of these values in the clan was relatively stable, reflected by similar distributions in consecutive years.

**Figure SI-2:**
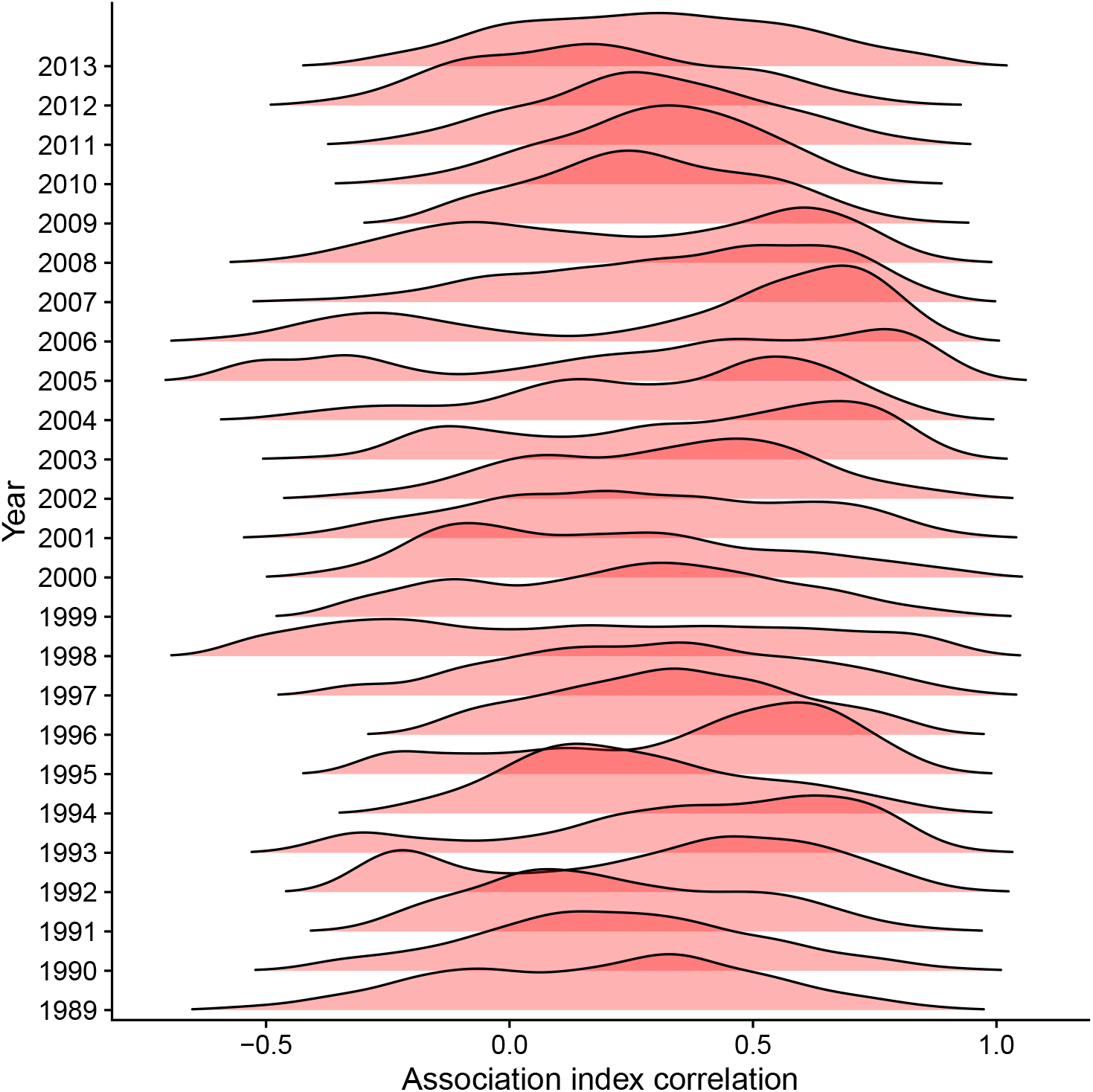
Distribution of correlation in association indices in spotted hyenas. Correlation in association index measures the similarity in association with others between two hyenas over one calendar year. Shading is used to show overlapping distributions. Between 1989 and 2015, the distribution of these values in the clan was not stable, reflected by dissimilar distributions in consecutive years.

**Figure SI-3:**
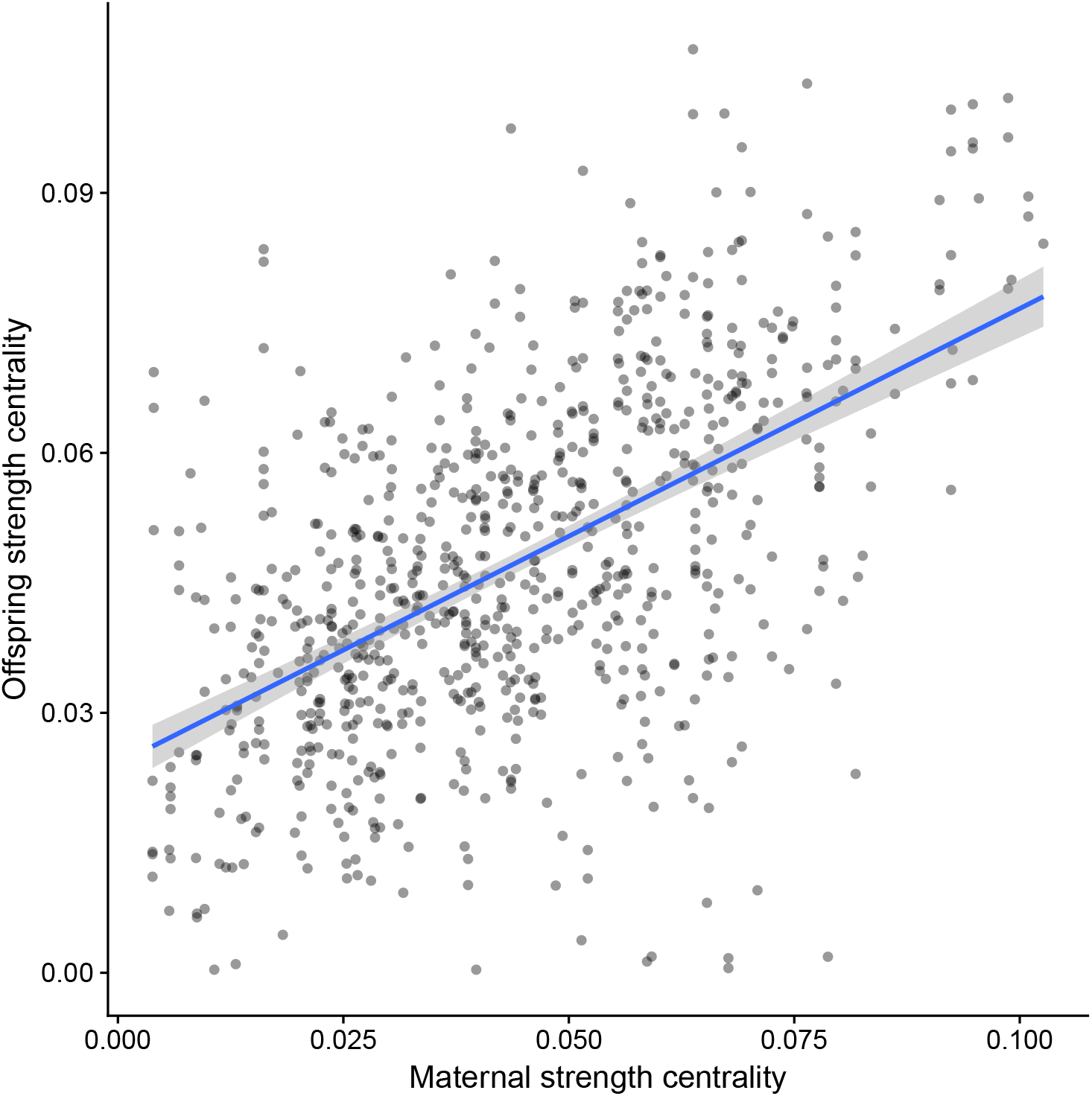
The centrality of mothers and their offspring. Strength centrality of offspring as a function of their mothers. Line depicts linear model and confidence intervals. *N* = 941.

**Figure SI-4:**
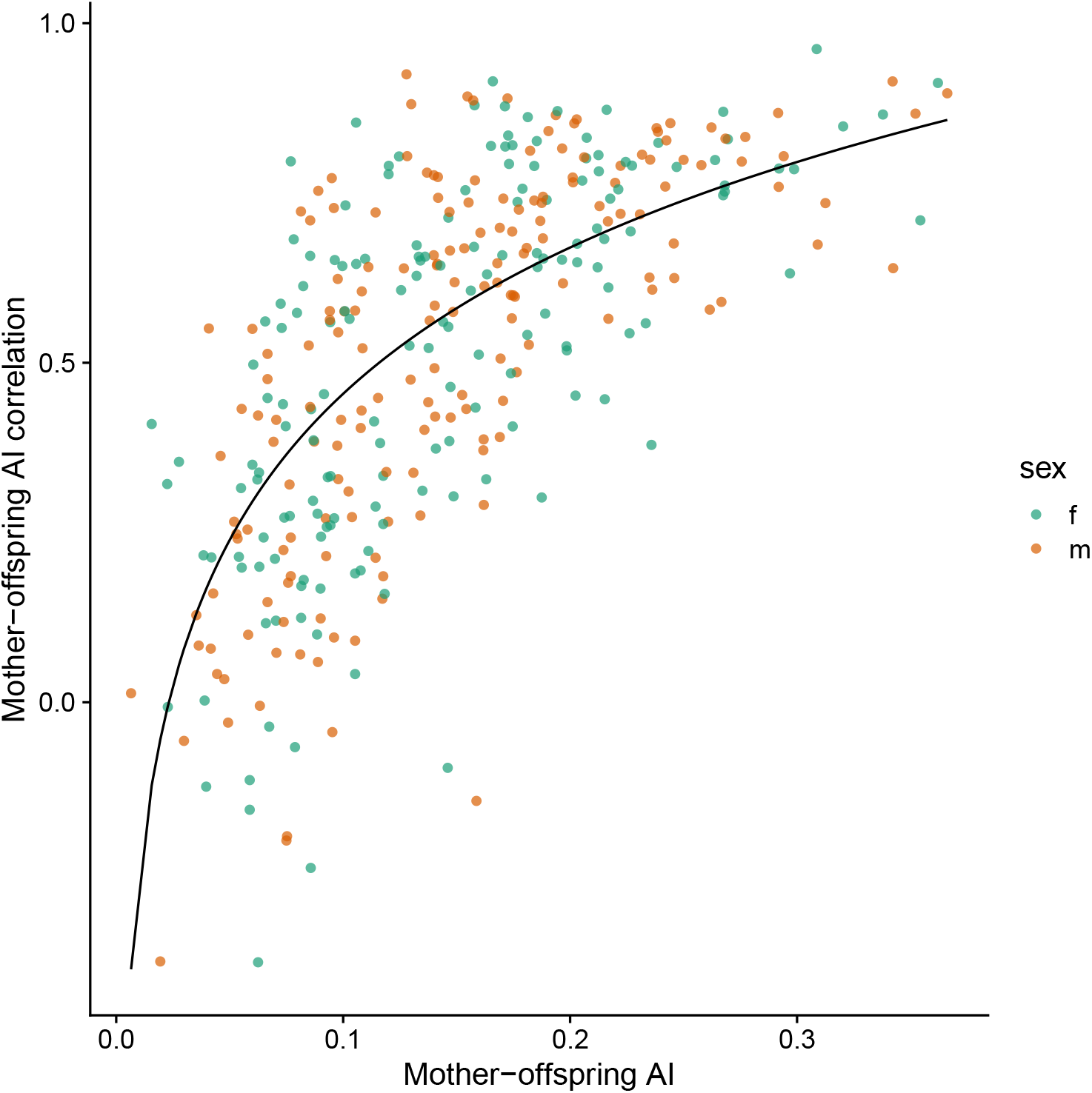
The effect of mother-offspring association on social inheritance. The strength of mother-offspring correlation in associations with others is plotted against mother-offspring association index for offspring in the first year after leaving the den. A curve is fitted using a GLMM model with mother identity and year set as random factors. The natural logarithm of Mother-offspring AI was used. Colors depict offspring sex. *N* = 347.

**Figure SI-5:**
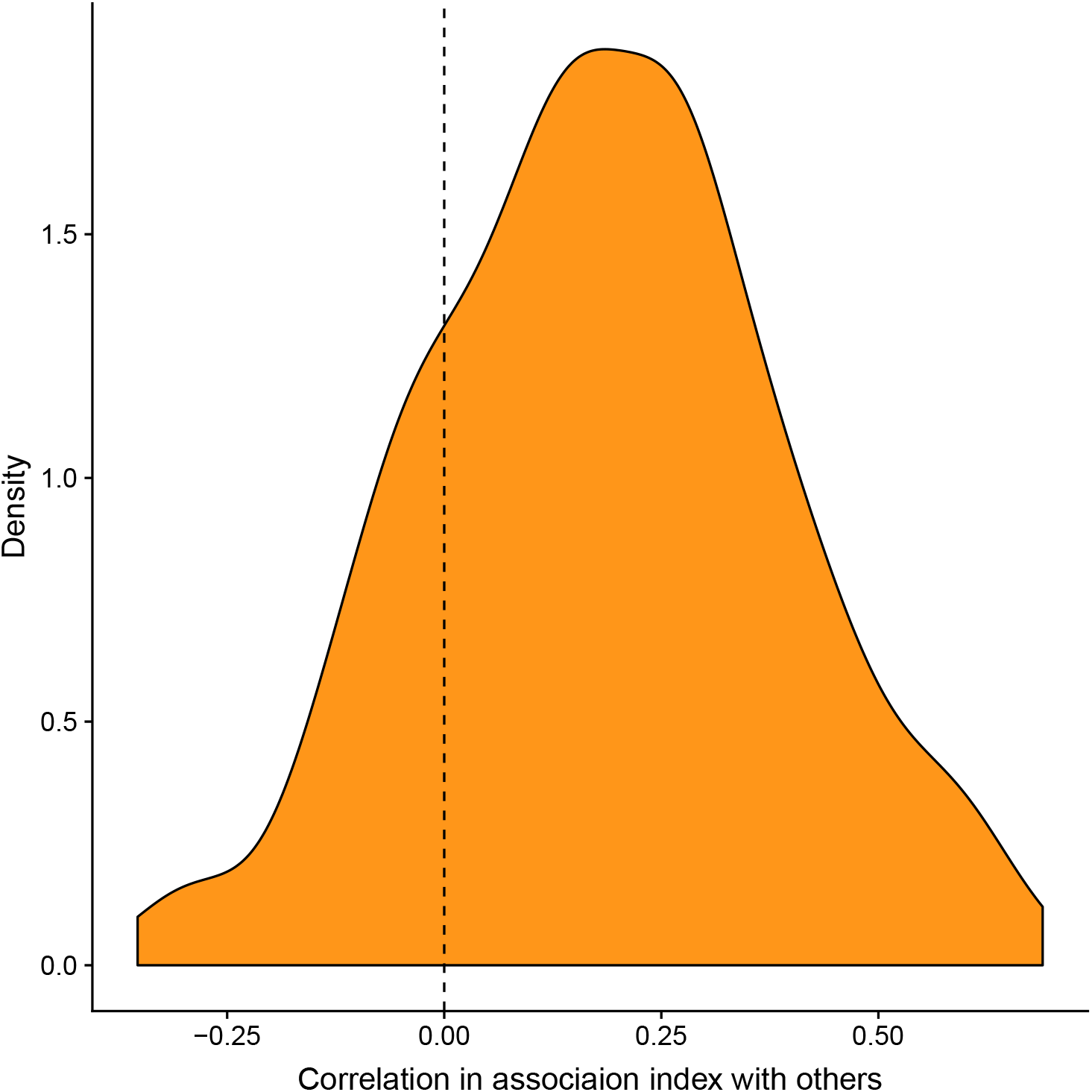
Correlation of associations across years. The distribution of correlations in AIs between hyena females and those of their offspring in the following year. Positive values indicate that offspring socially bonded with those who were previously associated with their mothers. The vertical dashed line depicts zero correlation. In a linear mixed model with mother identity as a random effect the intercept of mother-offspring correlation was 0.14 (*SE* = 0.03, *t* = 5.21)

